# Lactoferrin retargets adenoviruses to TLR4 to induce an abortive NLRP3-associated pyroptotic response in human dendritic cells

**DOI:** 10.1101/2020.06.03.131664

**Authors:** Coraline Chéneau, Karsten Eichholz, Tuan Hiep Tran, Thi Thu Phuong Tran, Océane Paris, Corinne Henriquet, Martine Pugniere, Eric J Kremer

## Abstract

Despite decades of investigations, we still poorly grasp the immunogenicity of human adenovirus (HAdV)-based vaccines in humans. In this study, we explored the role of lactoferrin, which belong to the alarmin subset of antimicrobial peptides that provide immediate direct and indirect activity against a range of pathogens following a breach in tissue homeostasis. Lactoferrin is a globular, iron-sequestering, glycoprotein that can increase HAdV infection and maturation of antigen-presenting cells. However, the mechanism by which HAdV-lactoferrin complexes induce maturation is unknown. We show that lactoferrin redirects HAdVs from species B, C, and D to toll-like receptor 4 (TLR4) complexes on human mononuclear phagocyte. TLR4-mediated internalization induces an abortive NLRP3-associated pyroptotic response inducing pro-inflammatory cytokine release and disrupting plasma membrane integrity without cell death. These data impact our understanding of the immunogenicity of HAdV-based vaccines and may provide ways to increase their efficacy.

## Introduction

The gaps in our understanding of the immunogenicity versus efficacy of viral vector-based vaccines are notable. For example, how does the rapid recruitment of immune cells and release of danger-associated molecular patterns (DAMPs) following vector delivery influence vaccine efficacy? In this study, we addressed the impact of a host defense peptide/protein (HDP) on human adenovirus (HAdV)-based vaccines. HDPs, also known as antimicrobial peptides, are evolutionary conserved effector molecules of the innate immune system. HDPs can act directly via antibiotic-like properties against a broad array of infectious agents [1,2], or indirectly by promoting the activation and/or maturation of antigen-presenting cells. Many HDPs are produced by neutrophils and epithelial cells of skin, oral mucosa, and the gastrointestinal tract. The cytoplasmic content of neutrophils, which are among the first leukocytes to infiltrate pathogen-infected and vaccine-injected tissues, is ~20% HDPs [3]. The rapid delivery of HDPs acts as part of the first line responders to the disruption of tissue homeostasis [4]. Functionally, HDPs are able to neutralize endotoxin, recruit and modulate the activities of immune cells, and induce angiogenesis. The alarmins (e.g. lactoferrin, α-defensin, and cathelicidin LL-37) are a subset of HDPs that also modulate innate and adaptive immune responses by directly engaging several pathways including pattern recognition receptor (PRR) signaling in antigen-presenting cells (APCs) [1,2]. Lactoferrin is an 80 kDa, multifunctional member of the transferrin family that sequesters iron, is produced largely by neutrophils, and its physiological concentration can reach mg/ml in some cases. Functionally, lactoferrin can induce dendritic cell (DC) maturation and, in the context of infections, drive Th1-cell responses [5–7].

In addition to their ability to influence innate and adaptive immune responses to bacteria, fungi, and enveloped viruses, some alarmins also influence adenovirus (AdV) uptake [8–10]. AdVs are 150 megaDaltons, ~90 nm diameter, nonenveloped proteinaceous particles containing a linear double-stranded DNA genome of ~36,000 (± 9,000) bp. Human AdVs (HAdVs) are classified into species (A-G) and types (~80) based on serology and phylogeny. In most cases, HAdVs cause self-limiting respiratory, ocular and gastro-intestinal tract infections in all populations regardless of health standards. Over the last 40 years the vectorization and immunogenicity of HAdVs have been of increasing interest in the context of vaccines, gene transfer, and morbidity associated with HAdV reactivation in immune-compromised individuals. In epithelial cells, alarmins influence HAdV infections via multiple mechanisms. Lactoferrin acts as a bridging factor during species C HAdV (types 1, 2, 5 and 6) infection in epithelial-like cells, independent of coxsackievirus adenovirus receptor (CAR), the primary cell surface attachment molecule for species C HAdVs [11]. Adams et al. reported that lactoferrin also mediates a modest increase in HAdV type 5 (HAdV-C5) uptake by human DC [9] and increased maturation. However, a mechanistic understanding of how increased uptake occurs and how DC maturation is induced, including which PRRs are engaged, are indispensable.

In this study, we characterize the mechanism by which HAdV-lactoferrin complexes induce human DC infection, inflammatory response, and maturation. We show that lactoferrin directly binds HAdV-C5, -D26, and -B35 with affinities in the micromolar range and increases HAdV uptake by mononuclear phagocytes. We demonstrate that lactoferrin re-targets HAdVs to toll-like receptor 4 (TLR4) complexes on the cell surface. Engagement of TLR4 complexes increases HAdV uptake, even in the presence of anti-HAdV neutralizing antibodies, and induces a cathepsin B-associated NLRP3 inflammasome, which includes caspase-1 activity, and release of interleukin 1 beta (IL-1β) - but not cell death. This pathway appears to be a variation of the alternative NLRP3 pathway induced by LPS via TLR4 engagement. In addition to a better understanding of the immunogenicity of HAdVs and HAdV-based vectors and vaccines, our data resolve the discordance between the TLR4-associated response to HAdVs in mice versus that of human phagocytes [12,13].

## Materials and Methods

### Cells and culture conditions

Blood samples were obtained from >100 anonymous donors at the regional blood bank (EFS, Montpellier, France). An internal review board approved the use of human blood samples. Monocyte-derived dendritic cells (DCs) were generated from freshly isolated or frozen CD14^+^ monocytes using CD14 MicroBeads human (MiltenyiBiotec) in the presence of 50 ng/ml granulocyte-macrophage colony-stimulating factor (GM-CSF) and of 20 ng/ml interleukin-4 (IL-4) (PeproTech). DCs stimulation was performed 6 days post-isolation of monocytes. Monocyte-derived Langerhans cells (LCs) were generated using 200 ng/ml GM-CSF and 10 ng/ml TGF-β. 911 cells and 293 E4-pIX cells were grown in Dulbecco’s modified Eagle medium (DMEM) and minimum essential medium (MEMα) with Earle’s salts, L-glutamine supplemented with 10% fetal bovine serum (FBS).

### Adenoviruses

The HAdV used in this study are replication-defective (deleted in the E1 region). The HAdV-C5 vector contained a GFP expression cassette. The HAdV-D26 vector contained a GFP-luciferase fusion expression cassette [14]. The HAdV-B35 vector contained a YFP expression cassette [15]. The vectors were propagated in 911 or 293 E4-pIX cells and purified to > 99% homogeneity by two CsCl density gradients.

### DC stimulation with HAdV-lactoferrin complexes

DCs (4 × 10^5^ in 400 μl of complete medium) were incubated with HAdV-C5, HAdV-D26 or HAdV-B35 (0.1 to 2 × 10^4^ physical particles (pp)/cell). We generated HAdV-lactoferrin complexes by incubating the virus with 40 μg lactoferrin (Sigma-Aldrich) for 30 min at room temperature. This corresponds to 100 μg/ml (1.25 μM) lactoferrin in 400 μl. These concentrations are similar to that found in an inflammatory environment of infected tissues. When specified, cells were complexed with IVIg (human IgG pooled from between 5,000 and 50,000 donors/batch) (Baxter SAS) or with lactoferricin (fragment of 49AA). Cells were incubated with HAdV-lactoferrin for 4 h, then washed and let incubated again for 24 h. The TLR4 agonist lipopolysaccharide (LPS) (Sigma-Aldrich) and NLRP3 inflammasome inducer nigericin (InvivoGen) were used at 100 ng/ml and 10 μM, respectively, to induce NLRP3 inflammasome formation. The inhibitors were used at the following concentrations, TLR4 inhibitors TAK-242 (Merck Millipore) at 1 μg/ml, oxPAPC (InvivoGen) at 30 μg/ml, TRIF inhibitory peptide (InvivoGen) at 25 μM, Syk inhibitor R406 (InvivoGen) at 5 μM, KCl (Sigma-Aldrich) at 45 mM, ROS inhibitor N-acetyl-L-cysteine (Sigma-Aldrich) at 2 mM, cathepsin B inhibitor MDL 28170 (Tocris Bioscience) at 0.1 μM, NLRP3 inhibitor MCC-950/CP-456773 (Sigma-Aldrich) at 10 μM, Bay11-7082 (Sigma-Aldrich) at 10 μM, caspase-1 inhibitor WEHD (Santa Cruz) and YVAD (InvivoGen) at 20 μM, VX765 (InvivoGen) at 10 μM, caspase-8 inhibitor Z-IEDT at 20 μM, RIPK1 inhibitor GSK963 (Sigma-Aldrich) at 3 μM, RIPK3 inhibitors GSK872 (Merck Millipore) at 3 μM and necrosulfonamide (R&D systems) at 1 μM. TLR4/MD-2, TLR4 (R&D Systems), MD-2 (PeproTech) recombinant protein and CD14 antibody (Beckman) were used at 20 μg/ml. Inhibitors were added on cells and recombinant proteins or antibody were added on HAdV-lactoferrin complex 1 h before stimulation. TLR4 surface expression level was assess with an anti-TLR4 antibody (Miltenyi Biotech) after 4 or 24 h.

### SPR analyses

Surface plasmon resonance (SPR) analyses were carried out on a BIAcore 3000 apparatus in HBS-EP buffer (10 mM HEPES, 150 mM NaCl, 3 mM EDTA, and 0.005% (v/v) polysorbate 20, pH 7.4). HAdV-C5, HAdV-D26 and HAdV-B35 diluted in acetate buffer at pH 4 were immobilized on three different flow cells of a CM5 sensor chip by amine coupling according to the manufacturer instructions. Immobilization levels were between 3,500 and 4,000 RU. Flow cell 1, without immobilized HAdV, was used as a control. Lactoferrin was injected at 100 nM on the four flow cells simultaneously. For K_D_ determination different concentrations of lactoferrin (6.25 - 200 nM) were injected at 30 μl/min during 180 s of association and 600 s of dissociation with running buffer. Regeneration was performed with pulses of gly-HCl pH 1.7. The kinetic constants were evaluated from the sensorgrams after double-blank subtraction with BIAevaluation software 3.2 (GE Healthcare) using a bivalent fitting model for lactoferrin. All experiments were repeated at least twice for each virus on a freshly coated flow cell.

### Flow cytometry

Cellular GFP or YFP expression from the HAdV-C5, -B35, -D26 vectors was assayed by flow cytometry. Fluorescence intensity was assessed after complex treatment for 24 h. Cell membrane integrity was assessed by collecting cells by centrifugation 800 × g, the cell pellets were re-suspended in PBS, 10% FBS, 7-aminoactinomycin D (7-AAD) (Becton-Dickinson Pharmigen) and analyzed on a FACS Canto II (Becton-Dickinson Pharmigen) or NovoCyte (ACEA Biosciences) flow cytometer.

Inflammasome formation was monitored as previously described [16] with minor modifications. DCs (1.5 × 10^5^ in 150 μl of complete medium) were seeded in a conical bottom 96 well plate and incubated with HDP-HAdV complexes containing 20,000 HAdV pp/cell. LPS/nigericin and immune complexed HAdV-C5 (IC-HAdV) were used as positive controls to identify inflammasome positive cells. IC-HAdV-C5 were prepared with IVIg (human IgG pooled from between 1,000 and 50,000 donors/batch) (Baxter SAS) as previously described [17]. Cells were fixed by adding 50 μl 4% PFA, PBS for 10 min on ice and centrifuged at 650 × g for 5 min. Supernatants were discarded and cells were permeabilized with 150 μl PBS/3% FCS/0.1% saponin for 20 min and collected by centrifugation. Supernatant was removed, and cells were re-suspended in 100 μl 1:500 rabbit anti-ASC (N-15)-R (Santa Cruz, sc-22514-R) PBS/3%FCS/0.1% saponin and incubated overnight at 4°C. Following overnight incubation, cells were pelleted at 650 × g for 5 min, washed once with 150 μl PBS/3% FCS:0.1% saponin, pelleted again and incubated for 45 min in 100 μl 1:500 Alexa-488 1:500 donkey anti-rabbit PBS:3% FCS:0.1% saponin for 45 min at room temperature. Cells were collected again by centrifugation and re-suspended in 150 μl PBS/ 3% FCS/0.1% saponin. The BD FACS-Canto II was used for acquisition. Samples were gated on DC and any doublets were excluded using forward light scattering (FSC)-area versus FSC width. Inflammasome positive cells were identified in the green channel as FL1-width low, and FL1-height high.

### Cytokines secretion

Supernatants were collected after 4 or 24 h and the levels of TNF and IL-1β were quantified by ELISA using OptEIA human TNF ELISA Set (BD Biosciences) and human IL-1β/IL-1F2 DuoSet ELISA (R&D systems) following the manufacturer’s instructions. In addition, 22 cytokines were detected by Luminex on Bio-plex Magpix using Bio-plex human chemokine, cytokine kit (Bio-Rad) following the manufacturer’s instructions.

### LDH release

LDH release was quantified using an LDH Cytotoxicity Assay Kit (Thermo scientific) following the manufacturer’s instructions. Briefly, 5 × 10^5^ cells were cultures in 96-well plates, infected for 4 h, and 100 μl of supernatant were collected to assess LDH activity. Fresh reaction mixture (100 μl) was then added to each well, incubated at room temperature for 30 min, the reaction was stopped, and the absorbance was determined at 490 nm using a microplate reader (NanoQuant, Tecan).

### Quantification of mRNAs

The levels of human *TNF, NLRP3*, *CASP1* and *IL1B* mRNAs were analyzed using quantitative reverse transcription-PCR (qRT-PCR). Total RNAs were isolated from DCs using a High Pure RNA isolation kit (Roche). Reverse transcription was performed with a Superscript III first-strand synthesis system (Invitrogen, Life Technologies) using 300 ng of total RNA and random hexamers. The cDNA samples were diluted 1:10 in water and analyzed in triplicate using a LightCycler 480 detection system (Roche, Meylan, France). PCR conditions were 95°C for 5 min and 45 cycles of 95°C for 15 s, 65°C or 70°C for 15 s, and 72°C for 15 s, targeting the *GAPDH* (glyceraldehyde-3-phosphate dehydrogenase) mRNA as an internal standard. Primer sequences were as follows for *NLRP3* (5’-CCTCTC TGATGAGGCCCAAG-3’ (*NLRP3* forward) and 5’-GCAGCAAACTGGAAAGGAAG-3’ (*NLRP3* reverse)) at 65°C, *IL1B* (5’-AAACAGATGAAGTGCTCCTTCC-3’ (*IL1B* forward) and 5’-AAGATGAAGGGAAAGAAGGTGC-3’ (*IL1B* reverse) at 65°C, *GAPDH* (5’-ACAGTCCATGCCATCACTGCC-3’ (*GAPDH* forward) and 5’-GCCTGCTTCACCACCTTCTTG-3’ (*GAPDH* reverse) at 70°C. Relative gene expression levels of each respective gene were calculated using the threshold cycle (2^−ΔΔCT^) method and normalized to *GAPDH* [17].

## Results

### Lactoferrin binds to HAdV-C5, -D26 and -B35 and increases infection of DCs

At physiological pH, HAdV-C5, -D26 and -B35 have patches of negative surface charges on hexon that should potentiate cationic alarmin binding. We therefore quantified the affinity of human lactoferrin to each virus capsid by surface plasmon resonance (SPR). The HAdVs were immobilized on a CM5 sensor chip and then escalating doses of lactoferrin were injected over the sensor surfaces. We found that lactoferrin binds the three HAdVs with affinities (K_D_) that varied from 0.8 to 54 μM (**Figure 1A** **&** **B**, **and** **Supplemental Figure 1A**).

**Figure 1.**
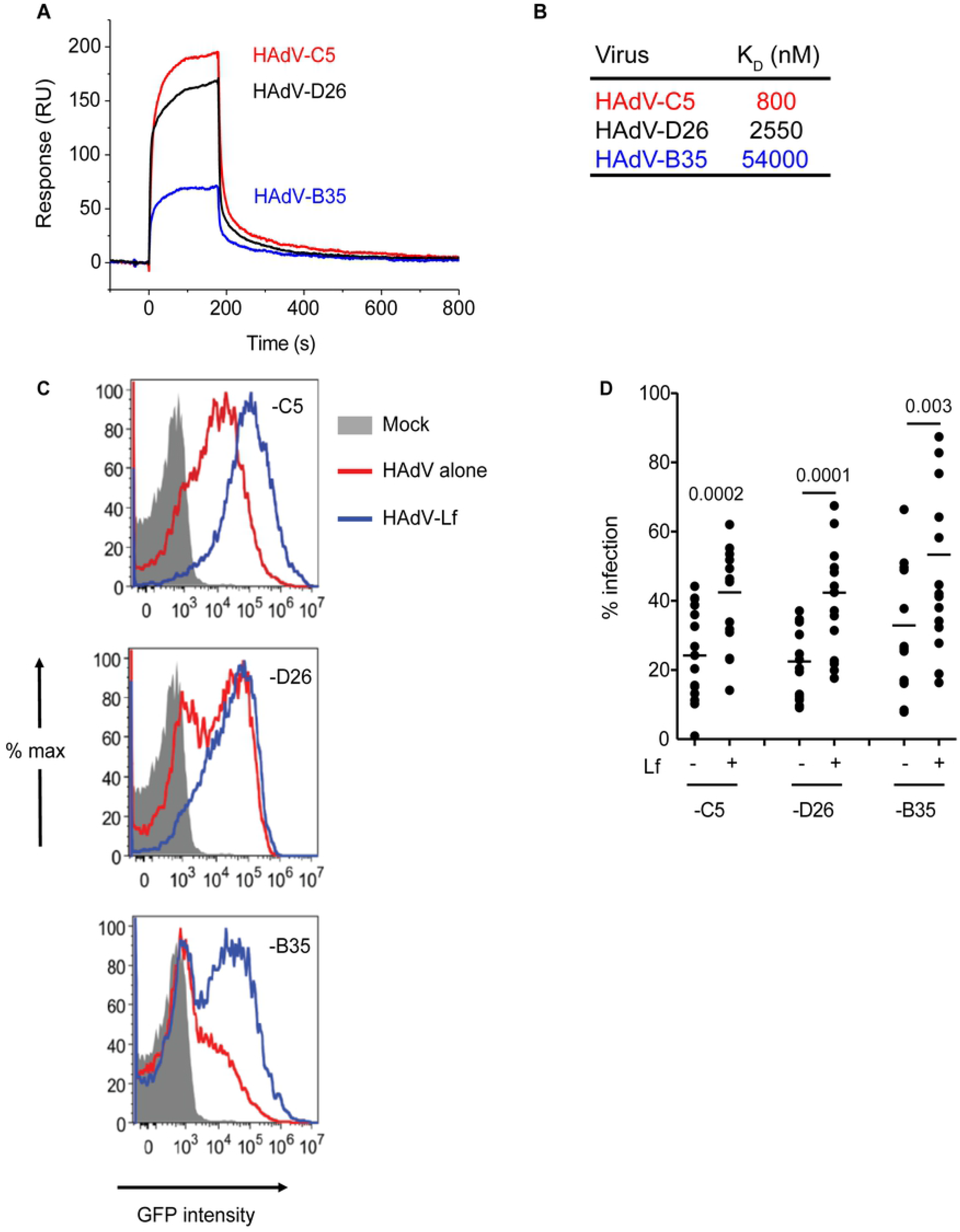
Lactoferrin binds to HAdVs and increase infectivity in DCs. **A)** Representative sensorgrams of lactoferrin binding to the HAdV capsid as assessed by surface plasmon resonance. HAdV-C5 (red), HAdV-D26 (black), and HAdV-B35 (blue) were covalently coupled to a CM5 sensor chip and lactoferrin was injected for binding comparison. For K_D_ determination a ranch of 6.25 - 200 nM of lactoferrin was injected and the K_D_ was calculated using a bivalent fitting model (RU = resonance units); **B)** Relative affinity (K_D_) of lactoferrin for HAdV-C5, -D26 and -B35 capsids; **C)** Representative flow cytometry profiles of DCs incubated with the HAdV vectors. DCs were mock-treated (grey) or incubated with HAdV species-C5 (5,000 pp/cell), -D26 (20,000 pp/cell) or -B35 (1,000 pp/cell) (red) or complexed with lactoferrin (blue) for 24 h. Samples were collected at 24 h, prepared for flow cytometry and 25,000 events were acquired/sample. **D)** Cumulative data (n = 21) from DCs generated from blood bank donors.

In human myeloid and epithelial cells, HAdV-C5, HAdV-D26, and HAdV-B35 receptor use does not overlap [18]: HAdV-C5 predominantly uses CAR as an attachment molecules, HAdV-D26 uses sialic acid-bearing glycans as a primary cell entry receptor [19], and HAdV-B35 predominantly uses CD46 [20]. We therefore tested the impact of lactoferrin on HAdV infection using replication-defective HAdV vector–mediated transgene expression, which is a surrogate assay for receptor engagement, internalization, cytoplasmic transport, docking at the nuclear pore, delivery of the genome to the nucleus, and transcription of the GFP expression cassette. Consistent with earlier reports [21], we found that lactoferrin increased (~2 to 4 fold) infection of human DCs by a species C HAdV (type 5), and in addition we show that HAdV-D26- and HAdV-B35-lactoferrin complexes are more infectious than the HAdV alone (**Figure 1C** **&** **D**). Preincubating HAdVs with lactoferrin, adding lactoferrin to the cell medium before HAdV, or adding lactoferrin to the cell medium after HAdV, all increased HAdV infection efficacy (**Supplemental Figure 1B)**. In addition, we found that lactoferrin-enhanced infection was not unique to DCs: -monocytes and monocyte-derived Langerhans cells were also more readily infected by HAdV-lactoferrin complexes (except HAdV-B35 on monocytes) (**Supplemental Figure 1C-D)**. Together, these data demonstrate that at physiological concentrations lactoferrin binds to three HAdV types from different species and increased uptake by DC.

### HAdV-lactoferrin complexes induce DC cytokine secretion

By influencing pathogen uptake alarmins could allow a host to better detect and respond to pathogens. Conversely, pathogen - alarmins interactions could reduce an APC’s ability to present antigens and therefore dampen a downstream response. If the response were pro-host, one would expect DC maturation and an inflammatory response. To determine whether HAdV-lactoferrin complexes influences DC maturation, we characterized the cytokine and chemokine profile using a multiplex array. Compared to mock-treated DCs, HAdV-D26 and HAdV-B35 induced a greater cytokine response than HAdV-C5 (**Figure 2A** **left column**). By contrast, compared to lactoferrin-treated DCs, HAdV-lactoferrin complexes induced an increase in the release of IL-1α and IL-1β (**Figure 2A** **center column**). When comparing HAdVs vs HAdV-lactoferrin complexes, the addition of lactoferrin induced a greater effect on HAdV-C5, which is likely due in part to the lower effect of HAdV-C5 alone (**Figure 2A** **right column**, **see** **Supplemental Figure 2A** **for raw data).** We then quantified IL-1β in the supernatant in time-dependent assays from multiple donors. HAdV-lactoferrin complexes rapidly induced >100-fold more IL-1β release than HAdVs alone, which further increased from 4 to 24 h post-stimulation (**Figure 2B)**. In addition, monocytes also released more IL-1β when challenged with HAdV-lactoferrin complexes compared to HAdV alone **(Supplemental Figure 2B)**.

**Figure 2.**
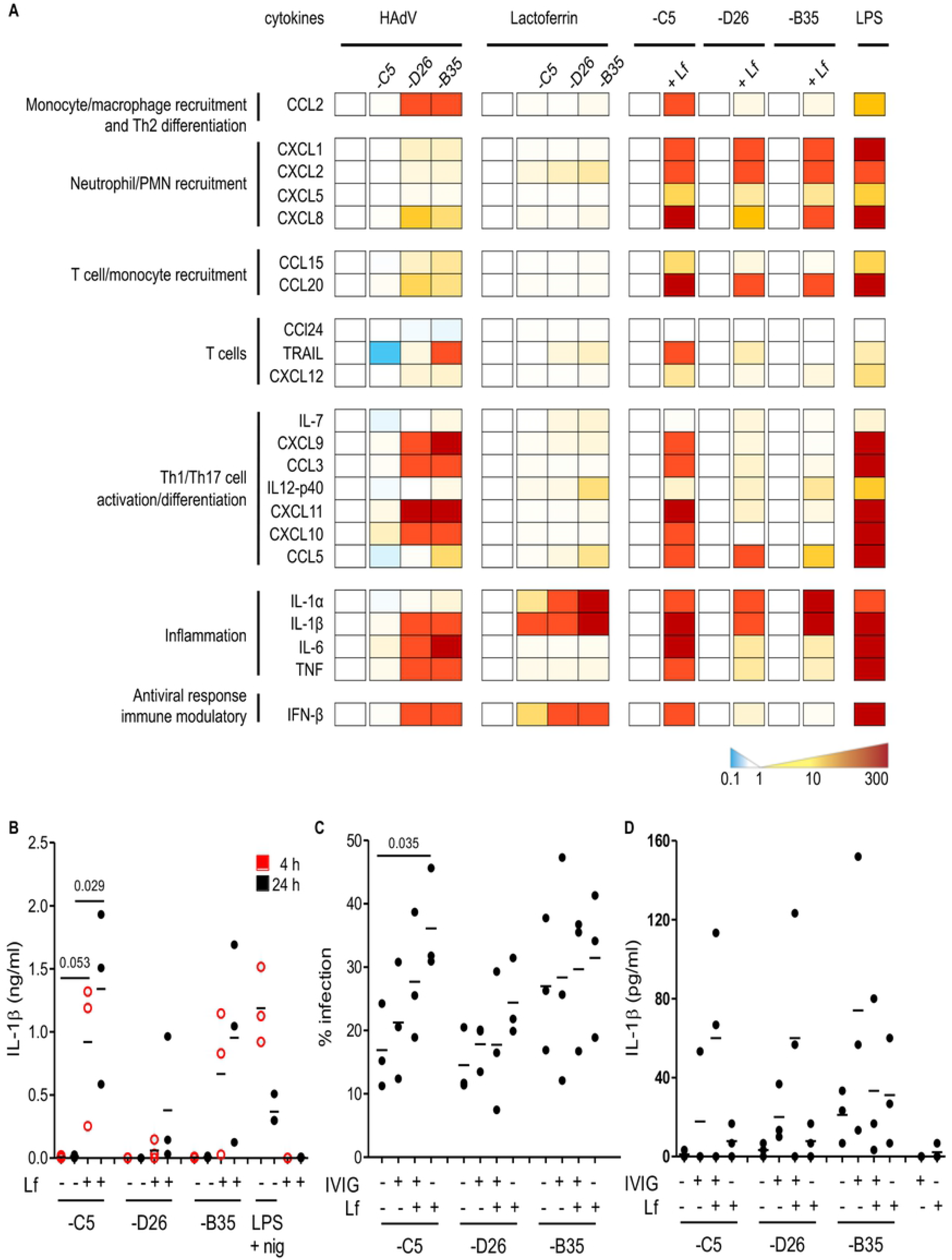
HAdV-lactoferrin complexes induce IL-1α and IL-1β. **A)** DCs were incubated with HAdV-C5, HAdV-D26, and HAdV-B35 ± lactoferrin (or LPS as a positive control) for 4 h and cytokines secretion in supernatants was assessed by Luminex. To the left of each set of columns is the baseline reference set: for the left-hand columns the baseline is mock-infected cells, for the middle columns the reference is lactoferrin-treated cells, for the right-hand columns the reference is HAdV-infected cells. Raw data can be found in (Supplemental Figure 2A & B). **B)** IL-1β release by DCs in the presence of HAdV-C5, -D26, and -B35 ± lactoferrin was assessed by ELISA at 4 (red) and 24 h (black) postinfection (n = 3). As control, cells were treated with LPS (TLR4 agonist) and nigericin (activated the inflammasome). **C-D)** DCs were incubated with HAdV complexed with lactoferrin, IVIG or lactoferrin + IVIG, for 4 h. Infection and IL-1β release, respectively, were analyzed 24 h postinfection (n = 3). Statistical analyses by two-tailed Mann-Whitney test.

We previously showed that individual serum and pooled IgGs (IVIG) containing anti-HAdV-C5 neutralizing IgGs induced the maturation of human DCs [17,22]. To benchmark the effects induced by lactoferrin, we compared HAdVs complexed with IVIG, lactoferrin or IVIG/lactoferrin. We found that HAdV-lactoferrin complexes led to a greater efficacy of gene transfer (higher percentage of cells expressing GFP). HAdVs complexed with IVIG/lactoferrin also led to more GFP/infected cell (more GFP^high^ cell) than HAdV-IVIG complexes (**Figure 2C** **and** **Supplemental Figure 2C**). Moreover, more IL-1β was released when IVIG and lactoferrin were combined with the HAdVs, consistent with observation that IL-1β release is directly linked to HAdV infection **(Figure 2D)**. In addition, HAdV-lactoferrin complexes decreased phagocytosis, a hallmark of functional DC maturation (**Supplemental Figure 2D)**. Together, these data demonstrate that lactoferrin-associated HAdV infection is associated with DC cytokine secretion and functional maturation.

### TLR4 is involved HAdV-lactoferrin induced DC maturation

Increased infection could be due to a handful factors, including alternative receptor engagement and/or more efficient intracellular trafficking. Of note, lactoferrin may increase DC maturation and IL-1β release by interacting with TLR4 [5,23–25]. In some myeloid cells TLR4 forms a complex with MD-2 for ligand binding [26] and with CD14 for TLR4 internalization [27]. MD-2 acts as a co-receptor for recognition of both exogenous ligands and endogenous ligands [26,28,29]. In addition, Doronin *et al.* proposed that HAdVs, via a murine coagulation FX-bridge, interact with murine TLR4 [12]. Yet, human FX did not act as a bridge for HAdV-C5 via TLR4 on human DCs [13]. We therefore probed the possible interactions between HAdV-lactoferrin complexes and the TLR4 pathway. To determine whether HAdV-lactoferrin complexes engage the TLR4 complex on the cell surface, we incubated the complexes with recombinant TLR4, MD-2, or TLR4/MD-2 dimers, or blocked CD14 on the cell surface with an anti-CD14 antibody. We found the greatest reduction of infection in the presence of the TL4/MD-2 dimer (**Figure 3A** **and** **Supplemental Figure 3A**). To address the involvement of the cytoplasmic TLR4 domain, we used TAK-242, a cell-permeable cyclohexene-carboxylate to disrupt TLR4 interaction with adaptor molecules TIRAP and TRAM [30–32]. We found that TAK-242 reduced HAdV-lactoferrin-mediated infection, IL-1β release and TNF secretion in DCs (**Figure 3B-C**). By contrast, interruption of TLR4-TIRAP/TRAM interactions had no notable impact on the infection of DCs, monocytes, or Langerhans cells by the HAdVs alone **(Supplemental Figure 3B - D)**. We then used oxPAPC and TRIF inhibitory peptide (Pepinh-TRIF) to inhibit extracellular TLR4 - MD2 interactions and cytoplasmic TLR4 - TRIF interactions, respectively. oxPAPC and Pepinh-TRIF decreased infection of HAdV-C5- and HAdV-D26-lactoferrin, while infection by HAdV-B35-lactoferrin increased (**Figure 3D-E)**.

**Figure 3.**
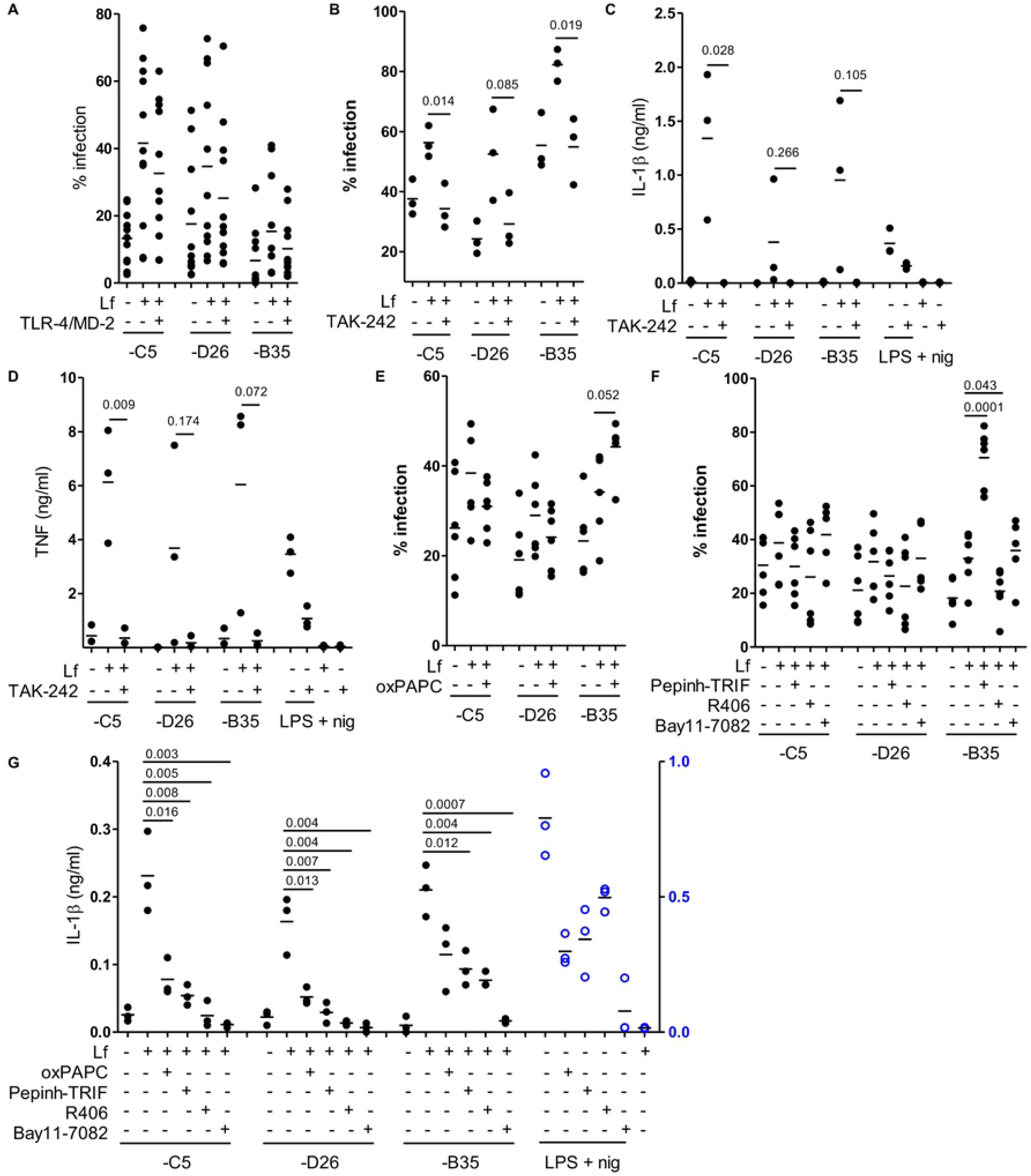
Blocking TLR4 engagement and signaling decrease HAdV-lactoferrin entry and DC maturation. **A)** HAdV-lactoferrin complexes were incubate with TLR4/MD-2 recombinant protein for 30 min. Infection was analyzed 24 h postinfection by flow cytometry (n =11); **B)** DCs were treated for 1 h pre-infection with TAK-242, HAdV-lactoferrin complex infection was analyzed 24 h postinfection by flow cytometry (n≥3). Cytokine profile of DCs treated HAdV-lactoferrin ± TAK-242; **C)** IL-1β release following inhibition with TAK-242; **D)** TNF levels 24 h postinfection ± TAK-242 (n ≥ 3); **E)** Percent infection following inhibition with oxPAPC (n = 6); **F)** Percent infection following inhibition with Pepinh-TRIF, R406 or Bay11-7082 (n = 6). **G)** IL-1β release from DCs incubated with HAdV-lactoferrin ± oxPAPC, Pepinh-TRIF, R406 or Bay11-7082 (n ≥ 3). Statistical analyses by two-tailed Mann-Whitney test.

We then perturbed TLR4-MyD88-Syk activation of the NF-κB pathway using R406 and Bay11-7082. R406 decreased HAdV-lactoferrin infections, consistent with its impact on TLR4-MyD88-Syk associated endocytosis, while Bay11-7082 had no effect on infection, consistent with its downstream signaling role (**Figure 3E)**. Importantly, all of the drugs and peptides affecting TLR4 interactions reduced IL-1β release (**Figure 3F**). In addition, lactoferrin, TAK-242, and Pepinh-TRIF did not change TLR4 surface expression, while high concentration of oxPAPC increased TLR4 levels **(Supplemental Figure 3E-G)**.

Of note, lactoferrin is also posttranslationally cleaved to generate lactoferricin, a biologically active N-terminal fragment of 49 aa. Lactoferricin also binds to negatively charged hexon hypervariable regions (HVRs) of HAdV-C5, -A31 and -B35 [33]. To determine if lactoferricin could mimic the effects of lactoferrin, we incubated the former with the HAdV vectors. We found no notable increase in HAdV infections or IL-1β release (**Supplemental Figure 3H** **&** **I**, respectively), suggesting that the C-terminal fragment plays a role in HAdV-TLR4 interactions. Together, these data demonstrate that interfering with TLR4 engagement/signaling reduces HAdV-lactoferrin-mediated transgene expression and DCs maturation in the case of HAdV-C5 and -D26.

### HAdV-lactoferrin complexes induce NLRP3 inflammasome formation

The inflammasome is a multiprotein cytosolic platform consisting of a PRR that induces nucleation of ASC (apoptosis associated speck-like protein containing a CARD), and recruitment of pro-caspase 1. Pro-caspase-1 auto-activation can be followed by removal of the N-terminal of gasdermin D (GSDMD), which initiates the loss of plasma membrane integrity via pore formation [34]. Classic NLRP3 inflammasome formation (canonical and non-canonical) is preceded by transcriptional priming event (signal 1) needed to produce inflammasome components and cytokines [35]. TLR4 engagement by LPS induces an alternative NLRP3 inflammasome activation, which does not need transcriptional priming, in human mononuclear phagocytes [36]. To determine whether HAdV-lactoferrin complexes induced transcription of inflammasome components, we used RT-qPCR to examine the mRNAs of inflammasome components. We found that in most cases HAdV-lactoferrin complexes significantly increased *NLRP3, CASP1, IL1B* and *TNF* mRNAs compared to the HAdVs (alone) (**Figure 4A**). To directly address inflammasome formation, we used flow cytometry to detect inflammasome-containing DCs using aggregation of ASC as a readout. During inflammasome formation, ASC changes from being distributed throughout the cytoplasm to an aggregate of ~1 μm diameter upon nucleation by a NLRP3. ASC nucleation can be directly visualized by changes in the fluorescence pulse width/ratio. While this assay does not allow quantification of all the cells that contain, or will contain an inflammasome, it does provide a snapshot of inflammasome formation at a given time. We found that <1% of mock-treated DCs contained an inflammasome. LPS/nigericin and HAdV-C5 complexed with neutralizing IgGs in IVIG (HAdV-C5-IgG) [17] had ~5 and 4% inflammasome-positive cells, respectively (~40% of the DCs will undergo pyroptosis in 8 h when incubated with this concentration of HAdV-C5-IgG [17]). We found an increase in the number of inflammasome-positive DCs 3 h post-challenge with HAdV-C5-lactoferrin complexes (**Figure 4B**). While the number of inflammasome containing cells were similar following challenges with HAdV-B35-lactoferrin complex compare to HAdV-C5- and HAdV-D26-lactoferrin, the difference was modest compared to HAdV-B35 alone.

**Figure 4.**
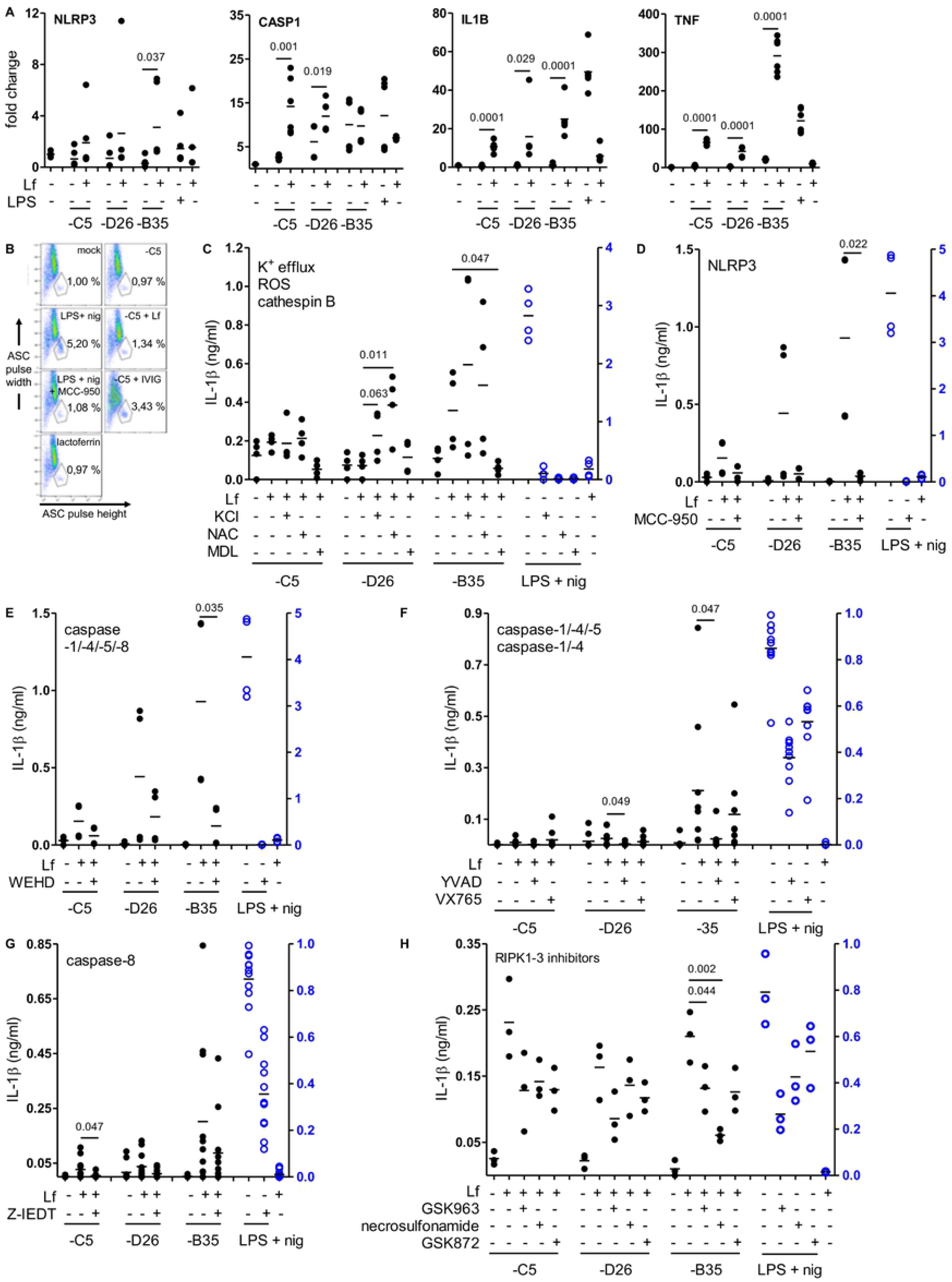
HAdV-lactoferrin complexes induce IL-1β via an NLRP3 inflammasome. **A)** mRNA expression of inflammasome components NLRP3 (NLRP3), caspase 1 (CASP1), pro-IL-1β (IL1B) and TNF (TNF) induced by HAdV-lactoferrin complex was assessed by RT-qPCR at 4 h postinfection; **B)** Flow cytometry-based assay for ASC aggregation (pyroptosome formation); Inflammasome formation was induced by incubating DCs with HAdV-C5-lactoferrin complexes for 3 h. LPS + nigericin and HAdV-C5-IVIG complexes were used as positive controls to identify inflammasome-positive cells. MCC-950 was used as a control of NLRP3 inhibition and lactoferrin on DCs was use as a negative control. Cells were stained with anti-ASC. Inflammasome-positive cells were identified as ASC-width low and ASC-height high. **C)** IL-1β release in response to HAdV-lactoferrin complex or LPS/nigericin in DCs pre-treated with NLRP3 pathway or caspase-1 inhibitor. IL-1β release in response to HAdV-lactoferrin complexes in DCs pre-treated with C) KCl, NAC, and MDL; D) NLRP3 inhibitor MCC-950; E-F) caspase-1 inhibitors WEDH, YVAD, VX765; G) caspase-8 inhibitor Z-IETD, H) RIPK1 inhibitor GSK963, and RIPK3 inhibitors necrosulfonamide and GSK872. N ≥ 3 in all assays. Statistical analyses by two-tailed Mann-Whitney test.

As IL-1β release is associated with both classic and alternative activation of NLRP3 inflammasomes, we explored the initiation steps. Inducers of NLRP3 inflammasome include K^+^ efflux, Ca^2+^ signaling, mitochondrial dysfunction, lysosomal rupture, or PRR engagement [34]. To identify the HAdV-lactoferrin-associated trigger(s), DCs were treated with KCl (to prevent K^+^ efflux), NAC (reactive oxygen species scavenger), and MDL (cathepsin B inhibitor). The addition of extracellular K^+^, and NAC did not decrease the release of IL-1β (**Figure 4C**). By contrast, MDL significantly reduced IL-1β release (**Figure 4C**), suggesting that rupture of lysosome-related organelles was involved. To determine whether the IL-1β release is linked to an NLRP3 inflammasome, DCs were pre-incubated with MCC-950 (NLRP3 inhibitor), which attenuated IL-1β release in response to HAdV-lactoferrin complexes (**Figure 4D**).

We then examined the role of the caspases by incubating cells with Z-WEHD-FMK (caspase-1, −4, −5 and −8 inhibitor), Z-YVAD-FMK (caspase-1, −4 and −5 inhibitor), VX765 (caspase-1 and -4 inhibitor) or Z-IETD (capsase-8 inhibitor). Globally, all inhibitors that affected caspase-1 and −8 activity reduced IL-1β release (**Figure 4E-G**). In addition to the direct effects of TLR4 engagement and signaling, it was possible that TNF secretion induced an autocrine response and inflammasome activation via the RIPK1-RIPK3-caspase-8 pathway [37]. While inhibition of the TNFR pathway (using GSK963, necrosulfonamide, and GSK872) had no significant effect on infection **(Figure 4H)**, IL-1β release was reduced in all cases. Possibly because HAdV-B35-lactoferrin complexes typically induced greater levels of IL-1β release, the effects of the caspase inhibitors tended to be more prominent **(Supplemental Figure 4A-C** for effect of inhibitors for each HAdV). Together, these data demonstrate that HAdV-lactoferrin complexes induce NLRP3 inflammasome formation and IL-1β release via a cathepsin B-mediated activation of caspase 1. Additionally, an autocrine effect of TNF may influence IL-1β release.

### IL-1β release without the loss of membrane integrity

In contrast to classic NLRP3 inflammasome activation, the alternative pathway does not include complete loss of cell membrane integrity (as based on LDH release into the extracellular space). This is thought to be due to ESCRT III pathway repairing pores in the plasma membrane induced by limited levels of GSDMD cleavage [38]. To determine if HAdV-lactoferrin complexes induce pores and the release of large intracellular proteins, we quantified extracellular levels of L-lactate dehydrogenase (LDH) activity at 4 h postinfection. LDH activity in the supernatant of control and HAdV-lactoferrin complexes was not significantly different from HAdV or lactoferrin-treated controls (**Figure 5A**). To determine whether HAdV-lactoferrin complexes were able to have a long-term impact DC membrane integrity we add a fluorescent marker of viability (7-AAD) to the DCs and quantified 7-AAD^+^ cells by flow cytometry. At 24 h post-challenge, the percentage of 7-AAD^+^ cell induced by HAdV-lactoferrin complexes was greater than lactoferrin- or HAdV-challenged cells (**Figure 5B**). Moreover, when lactoferrin was added to HAdV–IVIG complexes, we found an increase in the percentage of 7-AAD^+^ DCs (**Supplemental Figure 5)**.These data demonstrate that membrane integrity may be perturbed, but within the time frame of our assays cytosolic proteins are not releases into the medium. Together, these data demonstrated that HAdV-lactoferrin complexes prime DCs for NLRP3-associatedIL-1β release, inflammasome can be formed, membrane integrity is perturbed, but there is no significant leakage of cytoplasmic proteins.

**Figure 5.**
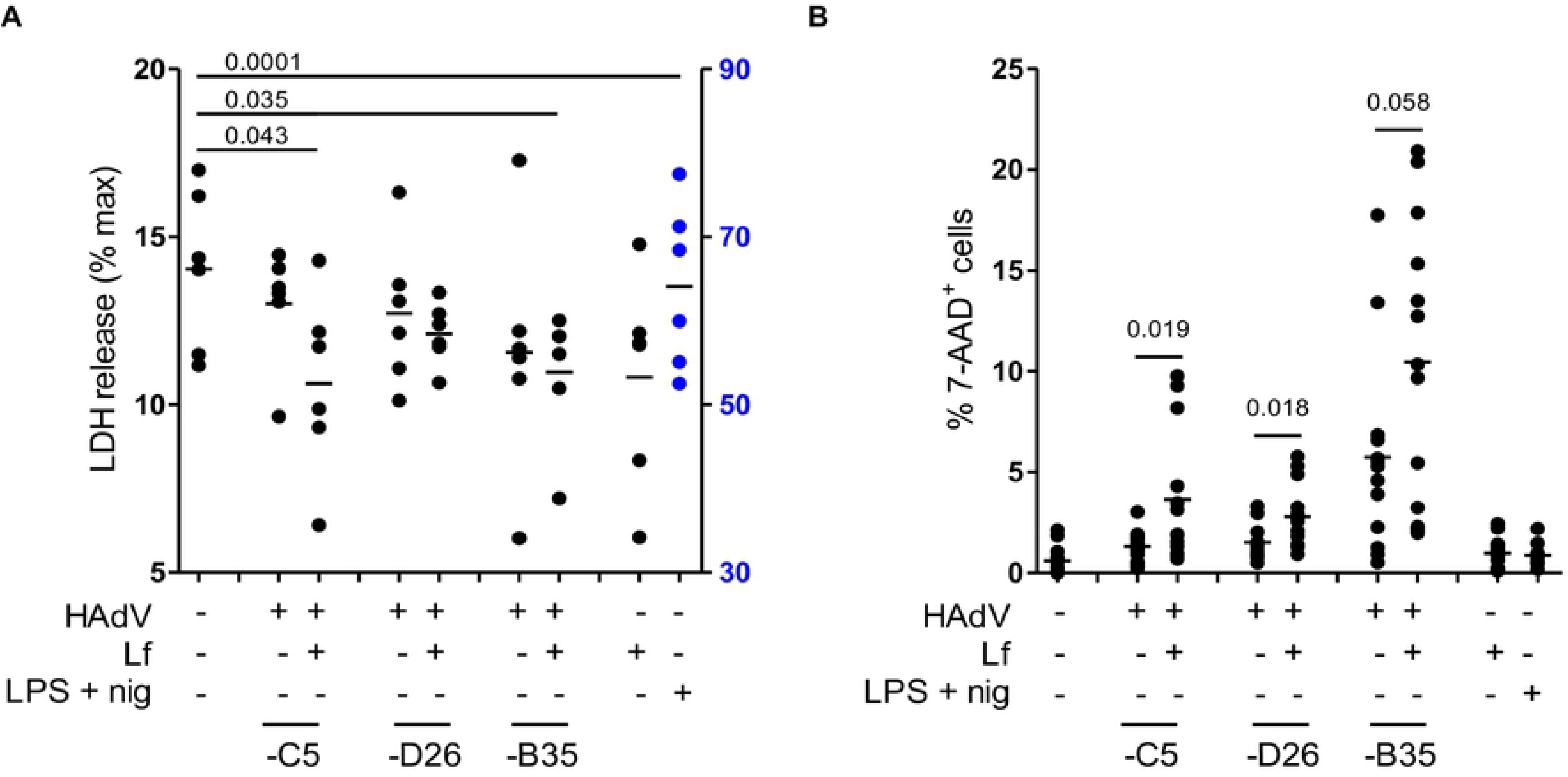
IL-1β release without the loss of membrane integrity. **A)** DCs were challenged with HAdV-lactoferrin complexes, HAdV alone, or LPS/nigericin. Loss of cytosolic content was quantified by LDH activity in the supernatant at 4 h postinfection (n = 6); **B)** Plasma membrane integrity, analyzed 24 h postinfection using 7-AAD uptake, was quantified using flow cytometry (n = 13). Statistical analyses by two-tailed Mann-Whitney test.

## Discussion

Deconstructionist approaches using binary systems to understand HAdV receptor engagement, trafficking, and immunogenicity provided a foundation to understand virus - cell interactions. Combinatorial assays using multiple human blood components can generate greater insight into clinically relevant HAdV issues, in particular tropism and immunogenicity. Here, we show how an alarmin influences the response of human DCs to three HAdV types. We examined pathways from receptor engagement, signaling, transcription, inflammasome formation/activation and cytokine release (**Figure 6**). Following engagement of TLR4, its TIR domain recruits MyD88 and TIRAP, which bridge TLRs to IRAK and MAPK family members that activate NF-κB, AP-1, and IRF. This latter pathway initiates transcription of genes coding for inflammasome components and proinflammatory cytokines [39,40]. The TIR domain also recruits TRAM and TRIF to activate the kinases TBK1 and IKK∊ to promote type I IFN expression [30]. Together, these innate immune pathways prime an adaptive antiviral response.

**Figure 6.**
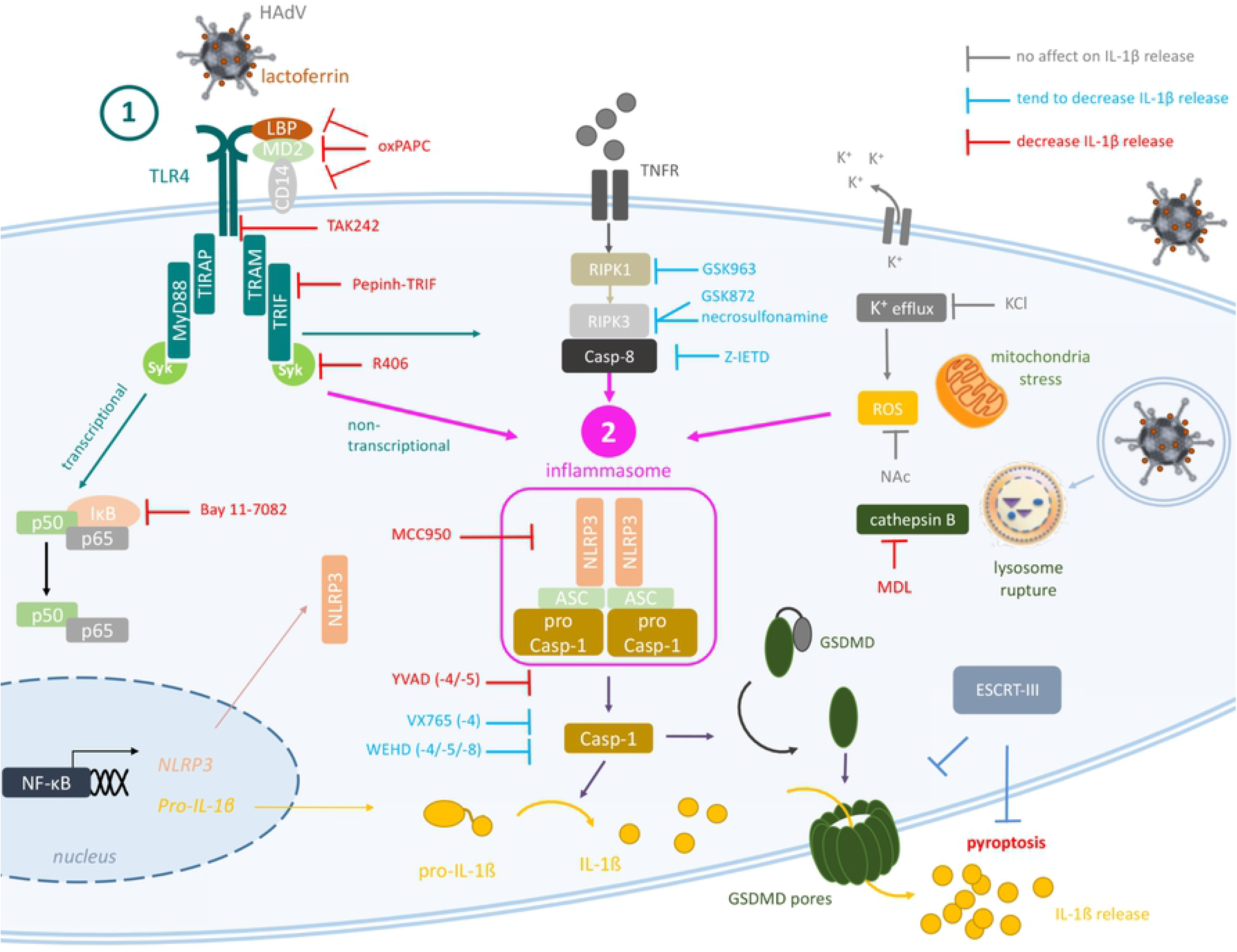
TLR4-mediated HAdV-lactoferrin uptake in DCs and IL-1β release. Lactoferrin binds to HAdV capsid and retargets the virus toward TLR4 complex on the cells surface. Following TLR4 engagement, its TIR domain recruits MyD88 and TIRAP, which bridge TLRs to IRAK and MAPK family members that activate NF-κB, AP-1, and IRF. This is a transcriptional priming event that initiating expression of genes coding for inflammasome components (e.g. NLRP3 and IL-1β). The DC then detects a second perturbations (signal 2) which induced ROS release (mitochondrial stress) or K^+^ efflux (perturbations of cellular integrity), and cathepsin B release from lysosome rupture. HAdV-lactoferrin complexes induce RIPK1/3 pathway through autocrine-TNF release or RIPK3 activation via TRIF. During inflammasome formation, pro-caspase-1 auto-activation induces cleavage of pro-IL-1β and likely GSDMD, which will initiate the loss of plasma membrane integrity via pore formation, allowing IL-1β release. Twenty-four hours post-challenge, DCs membrane integrity is intact, consistent with the involvement of ESCRT-III complex and repairing GSDMD pores.

Due in part to the technical advances in the synthesis of peptide-polymer conjugates, interest in new and old HDPs is flourishing. However, the complex biological functions of naturally occurring HDPs provide a candid reminder of how little we grasp their impact on immune responses to most pathogens. The three HAdV types used in this study were based on their state of development as vectors for vaccines [41,42]. HAdV-C5, -D26 and -B35 come from different HAdV species, and are associated with different vaccines efficacy. The *raison d’être* for the use of HAdV-D26 and -B35 is that their low seroprevalence (at least in Europeans and North Americans cohorts) and may circumvent some concerns associated with pre-existing HAdV humoral immunity [43]. It is worth noting that HAdV-B35 seroprevalence is typically rare – whether this is due to the lack of infection or lack of production of neutralizing antibodies to HAdV-B35 is currently unknown. Throughout this study HAdV-C5 and -D26 tended to have similar profiles in all assays. By contrast, there were several instances where HAdV-B35 was notably different. Whether these differences can be attributed to the use of CD46 or differences in the level between monocytes, DCs or Langerhans is possible, but unexplored. In addition, work in T cells that shows CD46 primes the NLRP3 inflammasome and therefore a possible binary engagement through TLR4 and CD46 could impact the response to HAdV-B35-lactoferrin complexes [44]. The breadth of the lactoferrin-enhanced infection of the three HAdVs suggest that the interactions are charge based because of the significant differences in the HVR sequences, which make up much of the surface area of HAdVs. Previous studies demonstrated that some HDPs attenuate HAdV infection of epithelial-like cells [8,15,45–48]. By contrast, our results are similar to Adams et al. [9] and demonstrated that lactoferrin-enhanced HAdV infection of human monocytes, DCs, and Langerhans cells. By delving deeper into these initial observations, our study sheds light onto the mechanisms by which an HDP connects HAdV infection of mucosal tissues, or during vaccination, to drive innate and adaptive immune responses. Mechanistically, it appears that lactoferrin reduces infection of epithelial cells and increases uptake into phagocytes, which provokes a pro-inflammatory and antiviral cytokine response. In combination with vaccination studies in paradigms of primary and recurrent infection, our study will help us understand how this pathway modifies adaptive immune responses against HAdV vectors and/or the transgene.

Classical NLRP3 inflammasome activation involves a two-step process: PRR-derived signal 1 to upregulate transcription of inflammasome components and NLRP3 posttranslational modification. NLRP3 then detects perturbations of cellular integrity associated with K^+^ efflux (signal 2). Consequently, Nek7–NLRP3 interaction leads to pyroptosome assembly and caspase-1-induced maturation of pro-IL-1β and pro-GSDMD. The alternative pathway consists of NLRP3-ASC-pro-caspase-1 signaling and IL-1β release without the loss of cytoplasmic content via GSDMD-induced pyroptosis. Yet, the alternative pathway delineated in this study is not an indisputable fit and likely reflects the variability between LPS and a HAdV. TLR4-mediated endocytosis, which is well characterized for LPS, depends on the homodimerization of TLR4. LPS, the quintessential TLR4 ligand, is extracted from gram^−^ bacteria by CD14, which then transfer it to MD-2, which interacts directly with TLR4. TLR4 dimerization is induced by the Lipid A region of LPS. Given the icosahedral shape and the size (~90 nm) of the HAdV-lactoferrin complex, one would expect that lactoferrin binds to multiple sites on the capsid and induced TLR4 dimerization directly, or possibly assemblage of multiple dimers. Dimerization is then associated with a CD14-dependent migration [49] to cholesterol-rich regions of the plasma membrane and endocytosis via a TLR4 ectodomain-dependent mechanism. While this picture appears partially consistent with the uptake HAdV particles, CD14 levels on monocytes-derived DCs are very low or absent, suggesting that migration to lipid rafts is via another pathway. The involvement of cathepsin B, a product of lysosomal rupture, is a key result. Most TLR4 agonists examined to date do not have complex intracellular processing. This is not the case for HAdVs. The endosomolytic activity of protein VI, an internal capsid protein, prevents the efficient degradation of the HAdVs in DCs by enabling the escape of HAdV capsid from endocytic vesicles/lysosomes into the cytoplasm [50,51]. However, this trafficking process causes the HAdV double-stranded DNA genome become accessible to AIM2 (absent in melanoma 2) and in turn the initiation of an inflammasome. The makeup and processing of TLR4-associated vs. Fc◻ receptor-associated endocytic vesicles is, to the best of our knowledge, unknown. From the data here, it appears that TLR4-associated endocytic vesicles fused to the cathepsin B-bearing lysosomal vesicles before the rupture of these vesicles causes the inception of an NLRP3 inflammasome. Of note, we did not detect an involvement of the cGAS pathway (the inhibitor RU.512 had no effect on HAdV-mediated transgene expression or IL-1β release, data not shown) suggesting that in our assays TLR4-mediated endocytosis was not associated with significant degradation of the HAdV capsid. These data are consistent with increased GFP expression. By inhibiting caspase 1, we showed that it is activated and involved in IL-1β release. This effect was specific for IL-1β, as TNF secretion did not decrease (**Supplemental Figure 6A)**. Furthermore, caspase inhibitors had no effect on HAdV-lactoferrin entry (**Supplemental Figure 6B)**. Caspase 1 cleavage of GSDMD abolishes its intra-molecular auto-inhibition and induces pore-like structures of ~15 nm in diameter in the plasma membrane to breakdown the ion gradients. Alternative inflammasome activation can be triggered by a unique signal. LPS sensing induces a TLR4-TRIF-RIPK1-FADD-CASP8 signaling axis, resulting in activation of NLRP3 by cleavage of an unknown caspase-8 substrate independently of K^+^ efflux. The alternative NLRP3 complex likely has a modified stoichiometry. Although caspase 1 becomes mature and cleaves IL-1β, pyroptosis is not induced and IL-1β release by an unconventional mechanism that functions independently of GSDMD. Why the inflammasome in HAdV-lactoferrin challenged cells does not lead to GSDMD-mediated release of cytoplasmic content may be due to the spatial and temporal signals the cell is receiving during the activation phase. In classic NLRP3 inflammasome activation, signal 1 is received well before signal 2 (NLRP3 engagement). In our assays, HAdV-lactoferrin induced signals are received immediately before the NLRP3 induction. Notably though, inhibition of the NF-κB pathway had a dramatic effect on IL-1β levels. The lack of coordination of transcriptional priming and de-ubiquitination of NLRP3 [52] may preclude pyroptosis, and favor an immune response with a longer duration and trafficking of DC to lymph node to induce an adaptive immune responses. Inflammasome activation is thought to be crucial for the induction of cellular and humoral immune responses in the context of vaccinations. However, whether the involvement of the HAdV-lactoferrin NLRP3 axis drives T-cell responses towards a Th1 or Th2 phenotype needs further analyses. By contrast, controlling excessive inflammatory response is necessary. In addition to the expression of IL-1β, we also found notable levels of IL-1α. It is also possible that the effects of IL-1α supersede or preclude pyroptosis because IL-1α can promote the expression of genes involved in cell survival [53].

Of note, our results may also resolve one of many conundrums associated with the differences between murine and human responses to HAdVs. If murine HDPs interact with murine coagulation factors to bind HAdVs and induce a TLR4-assocaiated pro-inflammatory response in the mouse liver [12], then the addition of ubiquitous HDPs into this picture could resolve the paradox. Whether HAdV-coagulation factor-HDP-HAdV complexes are produced following intravenous injection in mice has not been addressed. In conclusion, using combinatorial assays and primary human blood cells we detailed the multifaceted interactions between a PAMP (HAdV), a DAMP (lactoferrin), and PRRs (TLR4 & NLRP3) at the interface of innate and adaptive immunity in humans. These data directly address how the multiple layers of the innate and adaptive immune responses coordinate reactions to pathogens.

## Acknowledgments

We thank the imaging facility MRI, member of the national infrastructure France-BioImaging supported by the French National Research Agency (ANR-10-INBS-04, “Investments for the future”). We thank Katryn Stacey (CHU Montpellier) for help with the inflammasome detection by flow cytometry. We thank EKL members for constructive comments.

## Author contributions

Study design & conception: KE, EJK

Project direction: EJK

Performed experiments; CC, KE, HT, TTPT, OP, CH

Analyzed data: all authors

Wrote the manuscript: CC, KE, HT & EJK

Secured funding: EJK

## Data and materials availability

All materials can be obtained through an MTA.

## Supplementary Materials

**Figure S1:**
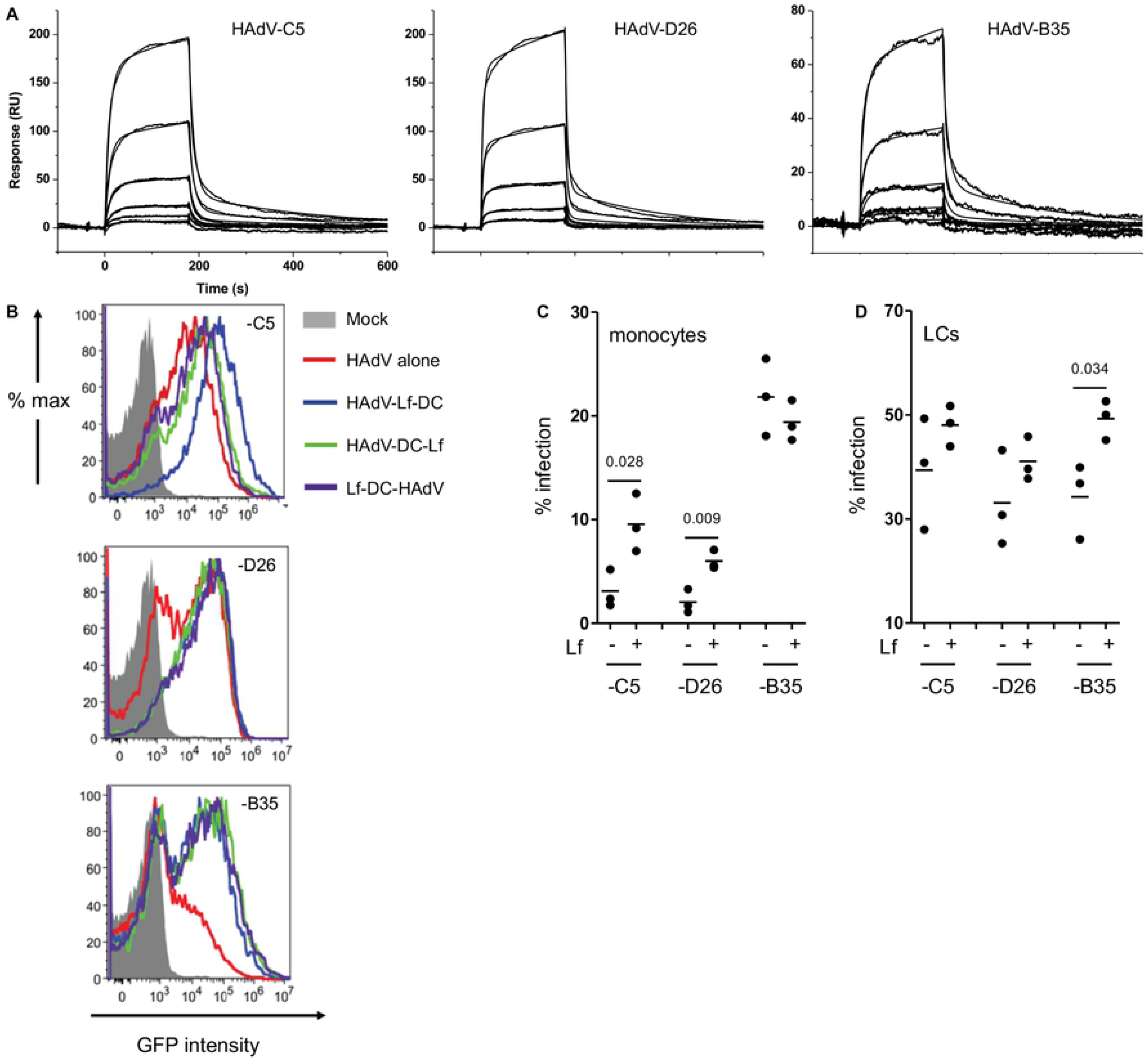
Analysis of lactoferrin binding to HAdV-C5, -D26 and −B35 by SPR. **A)** HAdV-C5, HAdV-D26, and HAdV-B35 were covalently couple to a CM5 sensor chip and escalating doses of lactoferrin (6.25-200 nM) for K_D_ determination. Depicted are overlaid sensorgrams (RU = resonance units); **B)** Representative flow cytometry profiles of cells infected with HAdV vectors. DCs were mock-treated (grey), incubated with HAdV species -C5, -D26 and -B35 alone (red), with lactoferrin complexed with HAdV (blue), with HAdV for 30 min and then lactoferrin (green) or with lactoferrin for 30 min and then HAdV (purple). Fluorescence was analyzed 24 h postinfection; **C)** monocytes and **D)** LCs were incubated with HAdVs ± lactoferrin and fluorescence was analyzed 24 h postinfection (n = 3).

**Figure S2:**
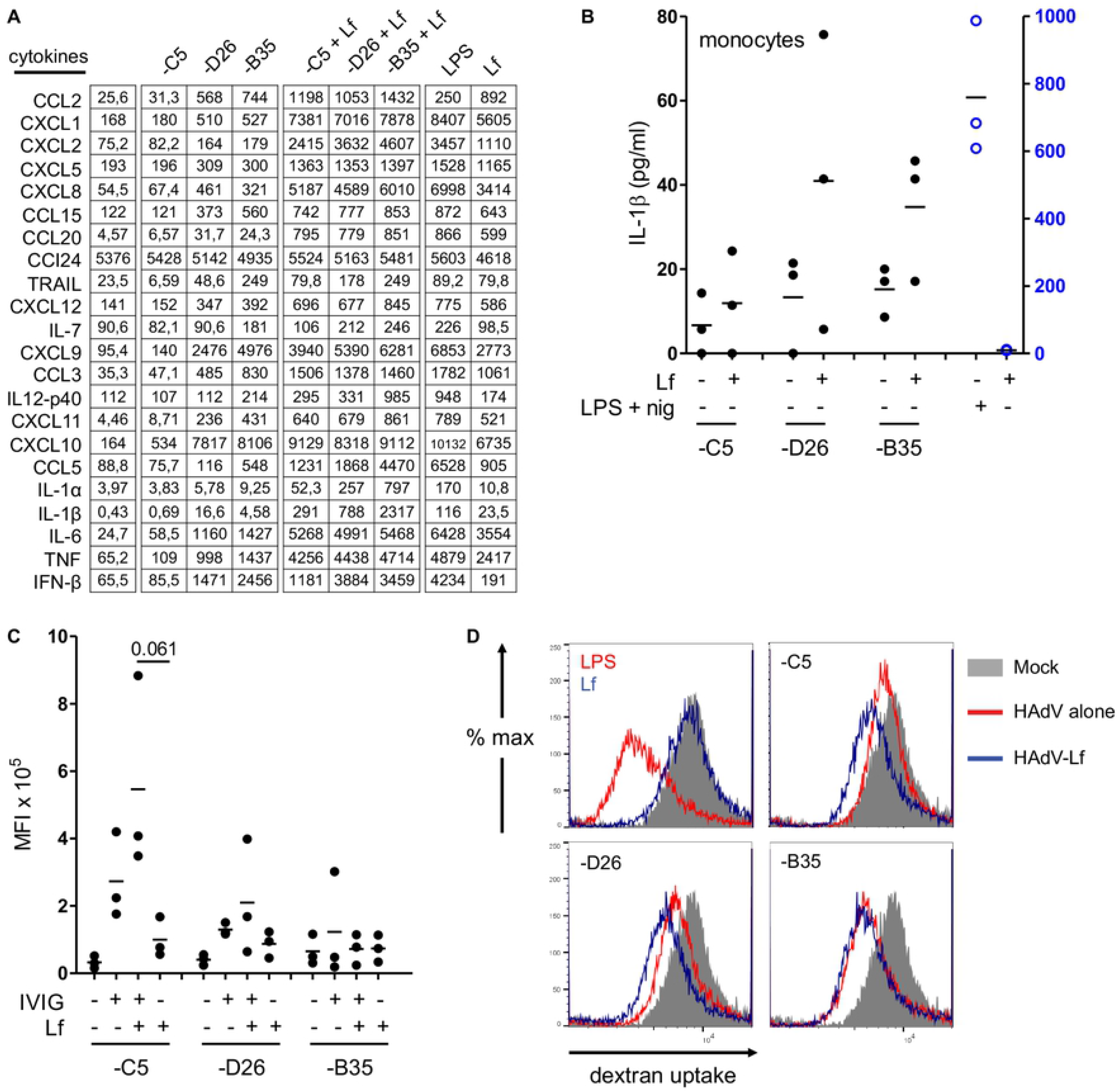
HAdV-lactoferrin complexes induce cytokine secretion. **A)** Raw data of Luminex assay of DCs ± HAdVs ± lactoferrin, plus controls. **B-C)** Freshly isolate human monocytes incubated with HAdV-C5-, HAdV-D26, and HAdV-B35 ± lactoferrin. The supernatants were used to quantify IL-1β release, GFP expression is reported median fluorescent index (MFI), respectively (n = 3) **D)** DCs were incubate with HAdV-lactoferrin complexes for 24 h. Cells were then incubated at 4°C or 37°C for 30 min with 1 mg/ml Texas Red-labelled dextran, washed with PBS, and immediately analyzed by flow cytometry to determine DC functional maturation (lower fluorescence = lower phagocytosis = greater maturation, n = 2).

**Figure S3:**
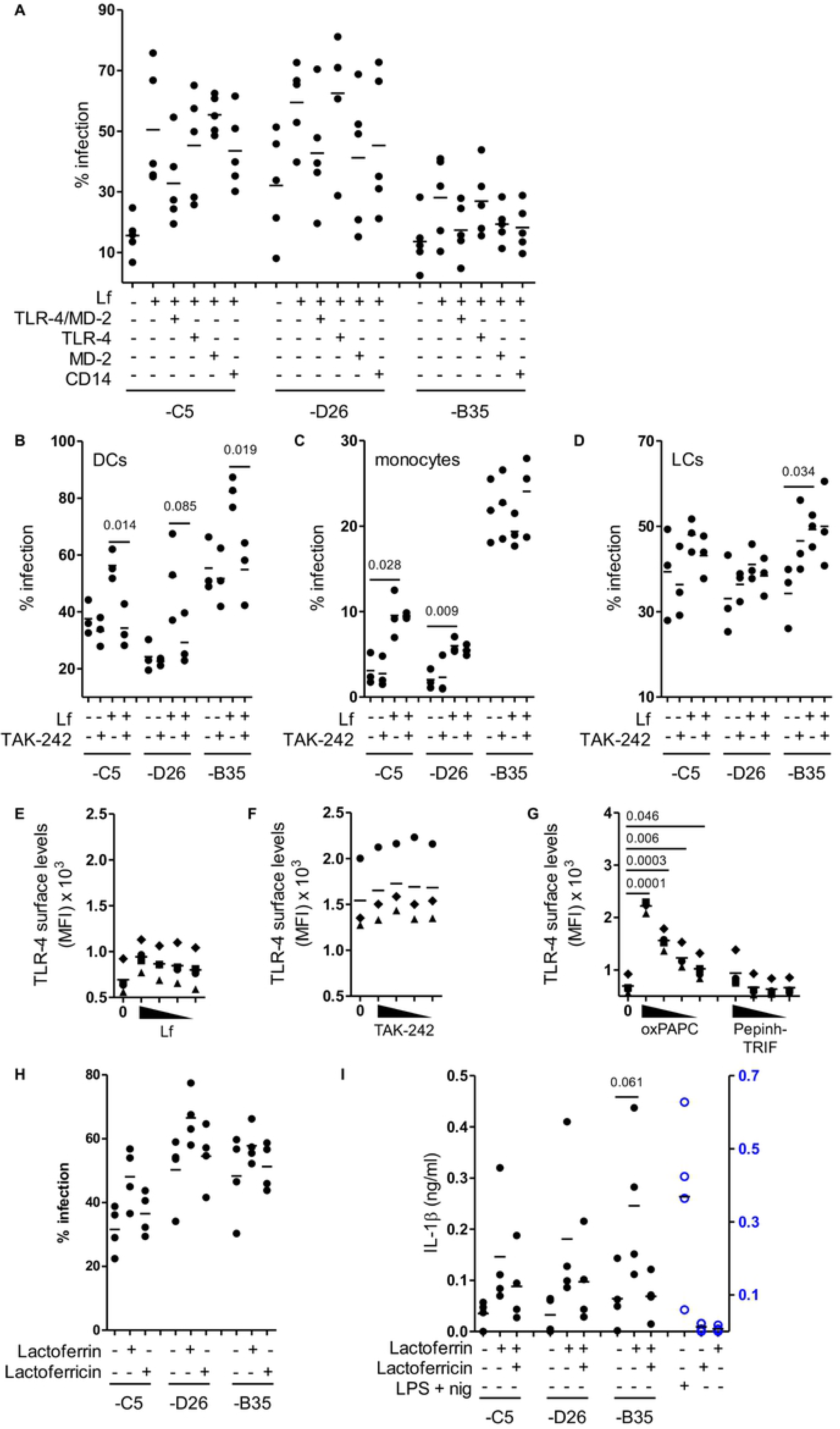
TLR4 inhibitors reduce infection of HAdV-lactoferrin complexes. **A)** HAdV-lactoferrin complexes were incubate with TLR4, TLR4/MD-2, MD-2 recombinant proteins or DCs were incubated with anti-CD14 antibody for 30 min. Then HAdV-lactoferrin-recombinant protein complexes were added to DCs or HAdV-lactoferrin were added to treated DCs. Infection was analyzed 24 h postinfection by flow cytometry (n = 4). **B-D)** DCs, monocytes, or LCs were treated for 1 h pre-infection with 1 μg/ml of TAK-242. HAdV-lactoferrin complex infection was analyzed 24 h postinfection by flow cytometry (n = 3). TLR4 surface expression was analyzed at 24 h after DCs treatment with decrease concentrations of **E)** lactoferrin (7 - 0.9 μg/ml), **F)** TAK-242 (200 - 25 μg/ml), **G)** oxPAPC (60 – 7.5 μg/ml) or Pepinh-TRIF (50 - 6.25 μg/ml) (n ≥ 3). HAdV were complexed with lactoferrin or with lactoferricin (a gift from H. Jenssen, Roskilde Universitet). **H-I)** DC infection and IL-1β release were analyzed at 24 h postinfection, respectively (n = 4). Statistical analyses by two-tailed Mann-Whitney test.

**Figure S4:**
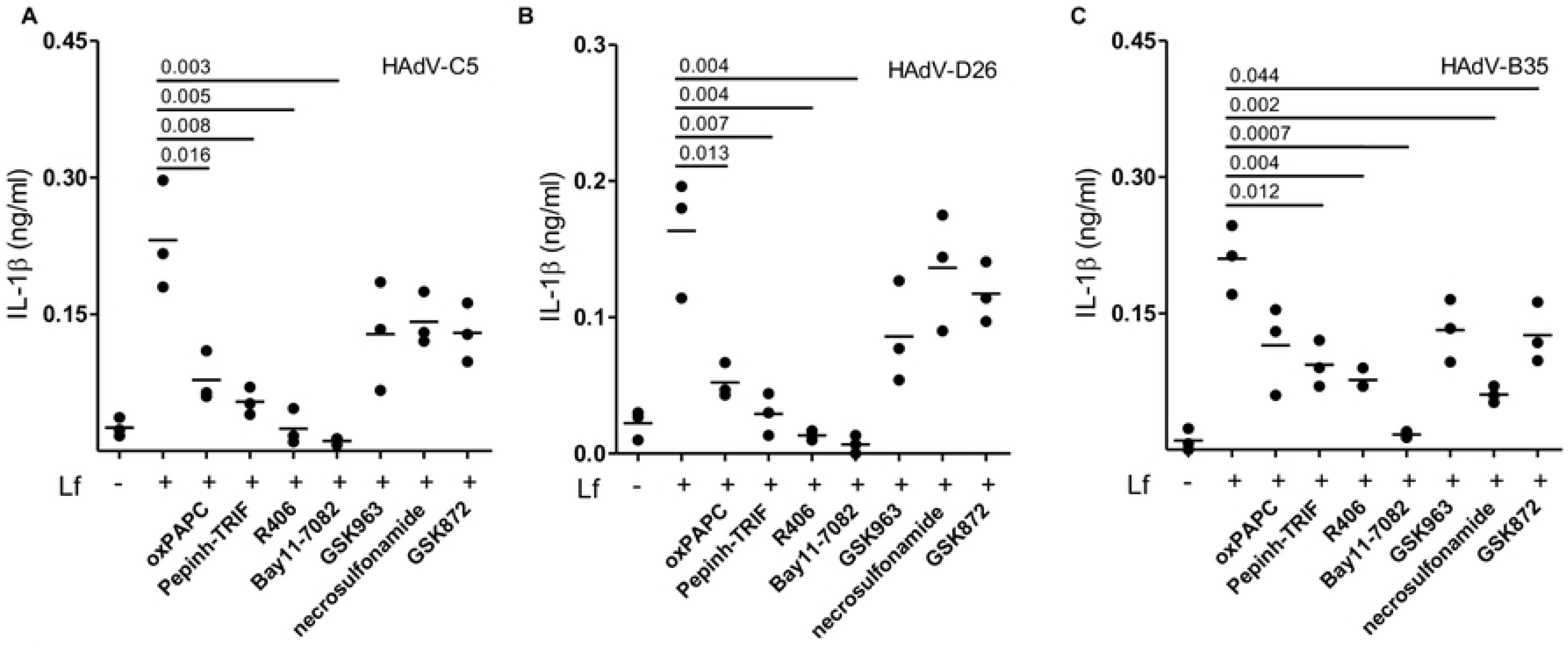
Pharmacological inhibition organized by HAdV type. DCs were treated for 1 h pre-infection with TLR4 inhibitors oxPAPC, Pepinh-TRIF, Syk inhibitor R406, NLRP3 inhibitor Bay11-7082, RIPK1 inhibitor GSK963 and RIPK3 inhibitors necrosulfonamide and GSK872 (n ≥ 3). Cells were infected with **A)** HAdV-C5-lactoferrin, **B)** HAdV-D26-lactoferrin or **C)** HAdV-B35-lactoferrin complexes and IL-1β release was analyzed 24 h postinfection (n ≥ 3). Statistical analyses by two-tailed Mann-Whitney test.

**Figure S5:**
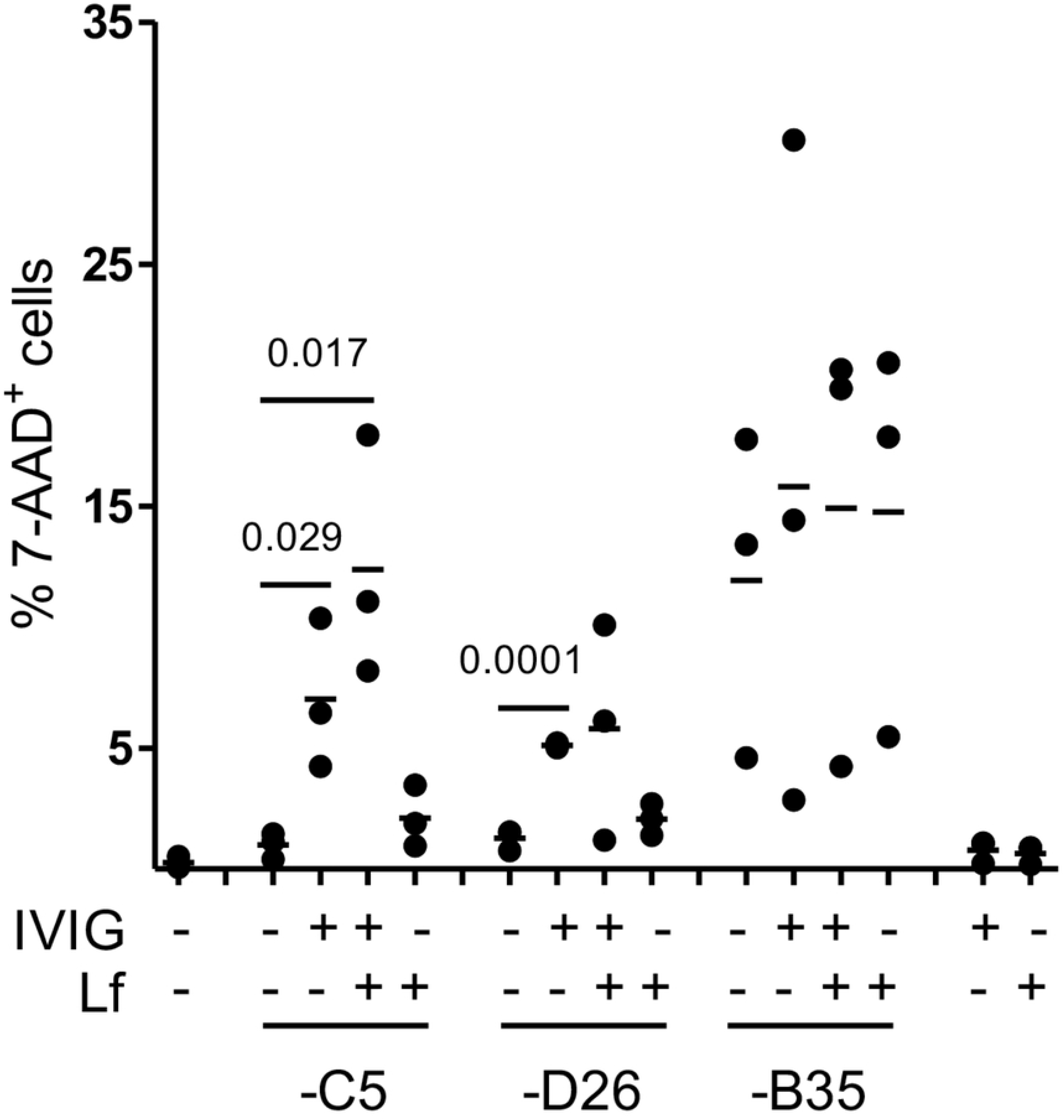
Lactoferrin combined with IVIG induced 7-AAD uptake. DCs were incubated with HAdV complexed with lactoferrin, IVIG or lactoferrin + IVIG. Plasma membrane integrity was analyzed by 7-AAD uptake by flow cytometry at 24 h postinfection (n = 3). Statistical analyses by two-tailed Mann-Whitney test.

**Figure S6:**
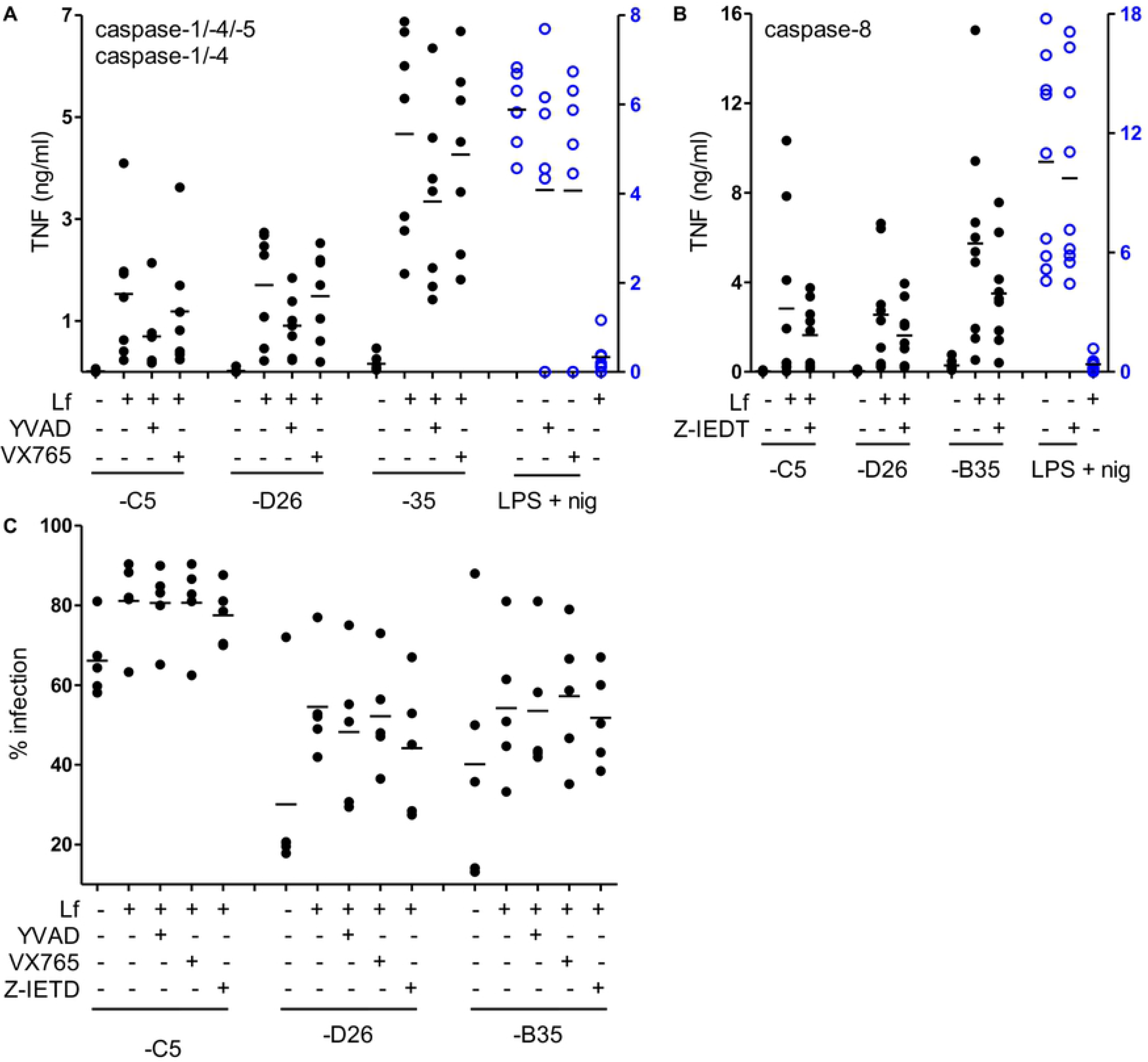
Caspase-1 and -8 inhibitors do not impact TNF secretion or infection. DCs were treat with caspase-1 inhibitors (YVAD or VX765) or caspase-8 inhibitor (Z-IETD) for 1 h. Cells were infected with HAdV-lactoferrin complexes for 24 h. **A-B)** Supernatant was collected for TNF quantification; and, **C)** cells for GFP expression (flow cytometry) (n ≥ 4). Statistical analyses by two-tailed Mann-Whitney test.

